# Scents for safe sex: a volatile sex pheromone promotes outbreeding in a parasitoid wasp with severe genetic load

**DOI:** 10.64898/2026.10.01.756042

**Authors:** Xavier Fauvergue, Karolina Pecharová, Inessa Buch, Alexandra Auguste, Nicolas Borowiec, Benoit Lapeyre

## Abstract

When mating among genetically related individuals results in inbreeding depression, various behaviors and life-history traits evolve to favor outbreeding. Long-range sexual communication following natal dispersal is one such adaptation. In parasitoid wasps with single-locus complementary sex determination (sl-CSD), inbreeding increases the production of unviable or sterile diploid males. Mastrus ridens is such a species, with a genetic load underpinned by sl-CSD, combined with a high potential for sibmating because siblings emerge in close proximity from the same host and become sexually active shortly after emergence. With this context in mind, we investigated the mate-finding strategy of M. ridens using complementary behavioral, chemical, and electrophysiological approaches. Wind-tunnel bioassays and field experiments showed that males orient toward virgin females and, to a lesser extent, mated females. This attraction from a distance appeared to be the sole form of conspecific attraction. Chemical analyses using static and dynamic headspace sampling and solvent extracts, followed by GC–MS, revealed several potential pheromonal candidates, including methyl 6-methylsalicylate. This compound elicited reproducible antennal responses in males in GC–EAD recordings. Altogether, our results suggest that mate finding in M. ridens is guided by a volatile sex pheromone emitted by females, with methyl 6-methylsalicylate as one putative component. Such mate-finding mechanisms promote outbreeding and may explain the low proportion of unfit diploid males observed in natural populations. Surprisingly, chemical analyses suggested that males also emit this compound, whereas neither males nor females showed behavioral responses to males. This apparent discrepancy suggests that attraction may depend on a more complex pheromonal signal and opens further avenues for investigation.

## Introduction

Mating among genetically-related individuals often comes with deleterious consequences on offspring survival and reproduction, a pervasive phenomenon referred to as inbreeding depression (Keller and Waller 2002). Inbreeding depression occurs because inbreeding increases individual homozygosity, revealing a pre-existing genetic load via the expression of deleterious recessive alleles or a reduction in the fitness benefits associated with heterozygosity (Charlesworth and Willis 2009). Although inbreeding tolerance is widespread (Szulkin et al. 2013, De Boer et al. 2021), a large body of empirical research shows that life history traits and behaviors that promote inbreeding avoidance have evolved repeatedly. Sibmating avoidance based on kin recognition (Krause et al. 2012, Leclaire et al. 2013, Galezo et al. 2022) and sex-biased dispersal (Shirane et al. 2019) have been extensively documented. Other mechanisms such as multiple mating (Tregenza and Wedell 2002), asynchronous maturation (Perrier et al. 2014), or reproductive suppression (Wolff 1992) were also interpreted as adaptations for inbreeding avoidance. Coupled with these mechanisms, outcrossing generally implies dispersal underpinned by efficient locomotor and cognitive skills allowing unrelated individuals to encounter one another away from the natal site. In this article, we study a gregarious parasitoid wasp in which sibmating leads to strong inbreeding depression, and provide converging lines of evidence — from behavioral, field, chemical, and electrophysiological data — that females emit a volatile sex pheromone that attracts males from a distance.

In insects, chemical communication is the privileged pathway for intraspecific interactions (Murlis et al. 1992, Ayasse et al. 2001, Symonds and Elgar 2008). Blends of chemical compounds are released by an emitter, and perceived by a receiver on which they elicit a specific behavioral or physiological response. These pheromones for intraspecific communication serve various roles, including aggregation and reproduction (Yew and Chung 2015). Aggregation pheromones typically attract conspecifics of both sexes and lead to the formation of groups around an emitter or a shared resource (Wertheim et al. 2005). In contrast, sex pheromones involved in mate finding generally elicit a sex-specific response, thereby facilitating encounters between potential mates. Other reproductive functions include mate choice, courtship and mating. In many insect species, long-range, volatile sex pheromones are true signals, the emission and reception of which have coevolved to facilitate mate encounter and are therefore beneficial to both males and females. Although the emission of sex pheromones by males has been documented (Landolt and Heath 1990, Ruther et al. 2007), females are most often the advertising sex (Wyatt 2014).

In parasitoid wasps, sex pheromones for mate finding have been evidenced in several tens of species via various approaches including field captures, behavioral experiments in laboratory setups, and chemical analyses (Ruther 2013, Böttinger and Stökl 2020, Tonga et al. 2026). These sex pheromones are generally emitted by virgin females (Fauvergue et al. 1999) but there are puzzling examples of species where females continue to attract males after mating despite refusing copulation attempts (Metzger et al. 2010b). In species such as *Nasonia vitripennis*, where mate search is restricted to the natal patch, males act as the advertising sex and emit pheromones whose titres correlate with sperm availability, allowing females to avoid sperm-depleted mates (Ruther et al. 2007, Ruther et al. 2009). Sex pheromones used for mate finding are sometimes non-volatile and laid while walking on foraging substrates (Fauvergue et al. 1995, Pompanon et al. 1997, Bernal and Luck 2007), but volatile sex pheromones are more common (Keeling et al. 2004, Ruther 2013). An interesting hypothesis is that pheromone volatility correlates with natal dispersal and outbreeding (Böttinger and Stökl 2020).

Long-range sex pheromones in parasitoid wasps are fascinating from an evolutionary perspective. First, in common with insects of the order Hymenoptera, parasitoids are haplodiploid and reproduce via arrhenotokous parthenogenesis: females develop from fertilized diploid eggs and males from unfertilized haploid eggs (Heimpel and de Boer 2008). Therefore, virgin females can produce parthenogenetic male offspring and sire grandchildren through the females mated by these males. For this reason, it has been hypothesized that compared to diploid organisms, females of haplodiploid species should invest less time and energy in sexual advertisement (Godfray 1990). Nevertheless, the production of both males and females, and hence the ability to find mates, should be advantageous once populations depart from infinite size and a balanced sex ratio (Verner 1965, Taylor and Sauer 1980, Fauvergue et al. 2008, Heimpel and de Boer 2008, Steiner and Ruther 2009). Second, many parasitoids are gregarious or quasi-gregarious, leading to a patchy spatial distribution that has long reinforced the view that local mating is the privileged pattern — a view further supported by extensive research on local mate competition (Hamilton 1967, Godfray 1994). However, meta-analyses and case studies have shown that, in parasitoids, local mating is more often the exception than the rule, suggesting that mate encounters typically occur away from the natal patch (Godfray and Hardy 1993, Hardy 1994). In this context, long-range sex pheromones are likely to play a key role in facilitating mate location (Fauvergue et al. 1999).

A third evolutionary perspective on long-range sex pheromones in parasitoids concerns their proximate function in facilitating mate location after natal dispersal, which may have evolved as an ultimate response to inbreeding risk. Haplodiploidy reduces the severity of inbreeding depression in parasitoids, as deleterious mutations are exposed in haploid males and efficiently purged from the population (Henter 2003). However, this purging mechanism does not apply to the complementary sex determination gene (*csd*), which is ancestral to Hymenoptera (Asplen et al. 2009) and widespread in parasitoids (Cook 1993, Van Wilgenburg et al. 2006, Heimpel and de Boer 2008). In species with single-locus complementary sex determination (sl-CSD), sex depends on both ploidy and heterozygosity at the *csd* locus: unfertilized eggs, hemizygous at this locus, develop into normal haploid males (Whiting 1943); fertilized eggs develop into females if heterozygous, but into diploid males if homozygous. These diploid males are generally unviable or sterile (Cowan and Stahlhut 2004, Heimpel and de Boer 2008, Harpur et al. 2012). Sl-CSD thus represents a specific form of inbreeding depression based on overdominance, where no single allele is intrinsically deleterious (Charlesworth and Willis 2009). Its consequences can be severe: in brother–sister matings, for example, up to 25% of fertilized eggs may develop into diploid males. In turn, this selects for behavioral traits that promote outcrossing (Ode et al. 1995, Ruf et al. 2011, Harpur et al. 2012, Thiel et al. 2013, Harper et al. 2016, Martínez et al. 2021).

Here, we focus on the parasitoid *Mastrus ridens* (Hymenoptera: Ichneumonidae), a species in which several life-history traits suggest that selection should have favored the evolution of inbreeding avoidance mechanisms. Sex determination in this species is consistent with single-locus complementary sex determination (Retamal et al. 2016). Although unfit diploid males have been reported at high frequencies in bottlenecked laboratory cultures (Retamal et al. 2016, Zaviezo et al. 2018, Sandanayaka et al. 2022), they are rare in field populations from the native range (Retamal et al. 2016), suggesting that mechanisms limiting inbreeding may operate in the wild. Immature stages develop gregariously on cocooned larvae of the codling moth Cydia pomonella (Lepidoptera: Tortricidae), a major global pest of apples, pears, and walnuts. Up to ten parasitoids may develop on a single host (Sandanayaka et al. 2011a), and both sexes become reproductively active within hours of emergence (Sandanayaka et al. 2011b). Development is slightly protandrous, with males emerging on average one to two days earlier than females (Sandanayaka et al. 2022). This modest protandry may reduce opportunities for sibmating without necessarily preventing temporal overlap among siblings. Moreover, no behavioral avoidance of kin has been observed to date (X. Fauvergue, unpublished data). Given the costs of inbreeding, the clustered emergence of sexually active siblings, and the apparent absence of kin discrimination, both natal dispersal and efficient mate location mechanisms are expected to evolve. In this study, we demonstrate that *M. ridens* females emit a volatile sex pheromone that attracts males from a distance. Among the volatile compounds detected, methyl 6-methylsalicylate emerged as a candidate pheromone component and elicited consistent electroantennographic responses in males. Male attraction to female-emitted volatiles is robust both in laboratory and field conditions, even under harsh environmental constraints.

## Material and Methods

### Insects

All experiments were conducted on a strain of *Mastrus ridens* maintained in our laboratory (INRAE, Sophia-Antipolis). This strain resulted from individuals that we collected in apple orchards in Kazakhstan in 2018, admixed with two strains also founded with individuals from Kazakhstan but maintained in Chile (Pontificia Universidad Católica de Chile) and New Zealand (Institute for Plant and Food Research). Parasitoids were reared in 25 × 25 × 25 cm BugDorm 4M2222® cages (MegaView Science Co., Ltd.) at 23 ± 1°C and a 16:8 light-dark photoperiod. Wasps were fed with water and honey *ad libitum*. To promote outbreeding, each new cage was founded every week with 20-30 adults of both sexes emerging from different cages. This procedure proved efficient, as suggested by the low proportion of males among diploid adults in two annual quality-control assessments conducted during the experimental period. Based on random samples of adults emerging from rearing cages and genotyped at microsatellite loci to infer ploidy (Retamal et al. 2016), these proportions were 0.04 [95% CI: 0.00–0.20] and 0.00 [95% CI: 0.00–0.09], respectively. After one day of parasitoid acclimatization and mating in the cage, hosts were exposed for 3 to 7 days.

Hosts for parasitoid rearing were diapausing cocooned larvae of the codling moth *Cydia pomonella*. They were produced by simulating winter conditions in controlled-environment chambers set at 8:16 light:dark photoperiod, 17 ± 1℃ temperature and 55 ± 10% relative humidity. From eggs to mature larvae, hosts were reared individually in tubes filled with artificial diet (agar: 1.6%, cornmeal NPP: 19.4%, ascorbic acid: 0.5%, benzoic acid: 0.2%, nipagin: 0.2%). Fourth instar larvae were transferred to a 96-well PCR plate (one larva per hole) where they could spin a cocoon and enter diapause. This provided a convenient method to handle multiple hosts and expose them to parasitoids.

In order to control the age and mating status of tested individuals, parasitoid nymphs were transferred from rearing plates to individual transparent boxes where they could complete their metamorphosis and emerge (1.5 × 1.5 × 2 cm). Each day, newly emerged adults were placed into rearing cages sorted by date of emergence and sex, and maintained under these conditions before experiments.

Wasps without visible deformities were randomly allocated to a given mating status: virgin vs. mated males (VM, MM) and virgin vs. mated females (VF, MF). Because male *M. ridens* mate repeatedly (Sandanayaka et al. 2011b), male mating status was controlled according to experimental use. Virgin males were used in behavioral assays, whereas mated males were also used, when mentioned, for producing mated females and in some chemical analyses. Individuals assigned to the mating treatment were randomly paired in transparent tubes for up to one hour until mating was observed either directly or from video recording. Copulation was considered successful when the male mounted the female for at least 20 seconds with characteristic antennal up-and-down movements at the end of the sequence (Sandanayaka et al. 2011b). Wasps were then maintained in cages (one cage for each type) until the tests, i.e., 48-58 h after copulation for mated individuals.

### Behavioral assays in the wind tunnel

This experiment aimed to document long-distance sexual communication in *M. ridens* by testing the in-flight orientation of individual wasps (receivers) in response to airborne cues from conspecifics (putative emitters). The 3 × 4 factorial design involved three types of receivers – virgin females (VF), mated females (MF), and virgin males (VM) – crossed with four types of odor sources: the same three categories of conspecifics and a no-insect control (CT). Receivers were tested individually, whereas each conspecific odor source consisted of five wasps. All tested individuals were 2–5 days old, with the majority being 3–4 days old. The order of the 12 factor combinations was randomized whenever possible, with occasional adjustments dictated by insect availability and the need to maintain a balanced design over time. The number of replicates ranged from 26 to 30 per factor combination, totaling 339 tests.

Note that although we use the neutral terms “emitters” and “receivers”, these terms are used operationally and do not imply active signaling behavior. Indeed, we observed no specific behavior in *M. ridens* suggesting the active emission of volatiles. For simplicity, we also refer to the flask containing the emitters as the “odor source”, without implying prior knowledge of the nature or chemical identity of the volatile compounds potentially released by virgin or mated males and females.

The wind tunnel was previously described by Lo Pinto et al. (2004), Metzger et al. (2010b), and Auguste and Fauvergue (2015); a schematic diagram of the apparatus and its use in this study is provided in Figure 1. The flight chamber (150 × 70 × 60 cm) was ventilated by a constant laminar airflow of approximately 16 cm·s⁻¹. A secondary airstream was delivered at 1 L·min⁻¹ through a conical flask (5 × 5 × 10 cm, 175 mL) containing the emitters and was then injected into the main airflow. This secondary stream therefore carried any volatiles released by the insects into the main airflow. Receivers were released on a takeoff platform positioned 45 cm downwind from the odor-source and at the same height as the outlet. Two artificial plants (30 × 35 × 35 cm) were placed next to the flask exhaust tube to provide a landing site for parasitoids. Three additional, smaller artificial plants were positioned between the landing site and the takeoff platform. Several pieces of colored paper (blue, green, orange, pink, yellow) were randomly distributed on the floor, ceiling, and walls of the flight chamber to serve as visual cues.

**Figure 1.**
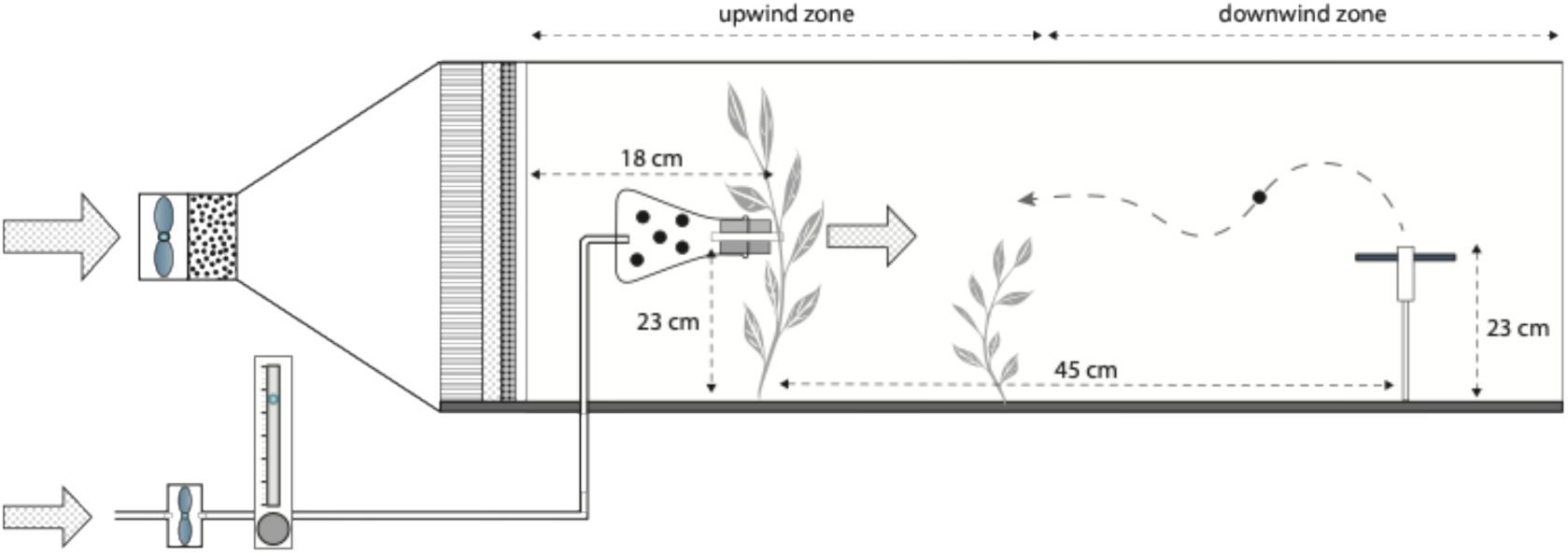
Schematic representation of the wind tunnel used to test flight responses in Mastrus ridens. A main laminar airflow traverses the flight chamber, combined with a secondary airflow that passes through a conical flask containing five individual emitters. The experiment followed a 3 × 4 factorial design, with both emitters and receivers varying in sex and mating status: virgin males, virgin females, and mated females, plus a control with no emitter. Conspecific individual receivers were released individually on a takeoff platform located 45 cm downwind. Their behavioral responses – takeoff, flight, and landing – were recorded as they navigated toward artificial plants positioned next to the flask.

Cages containing the insects to be tested were placed in the wind tunnel room at least 12 h prior to testing. Artificial lights were manually switched according to a regular 16:8 h light:dark schedule. All assays were conducted between 9:00 and 21:30, beginning at least 1 h after the lights had been turned on. Experiments were conducted at 24 ± 0.25 °C, 40–75% RH, and an uncontrolled atmospheric pressure ranging from 1007 to 1023 hPa.

On the day of testing, emitter flasks were prepared with five individuals of a given sex and mating status (virgin males, virgin females, or mated females), along with a small amount of water and honey as food. Control flasks were prepared identically, with water and honey but no insects. A flask of a given type was placed in the tunnel, and airflows were turned on. Prior to each test, an individual receiver wasp was allowed to acclimate for at least 5 min inside a tube positioned on the takeoff platform. A valid trial began when the wasp reached the opening of the release tube and became exposed to the odor plume. The trial lasted either five minutes or until the insect landed on the plant next to the source. If a parasitoid failed to leave the release tube within two minutes, it was removed from the tunnel and allowed to rest for at least five minutes before a new attempt. Individuals that failed to exit the release tube in three consecutive attempts were discarded. Each receiver therefore contributed to a single valid trial, although up to three release attempts were allowed before the individual was discarded. If a wasp completed the test without flying, its ability to fly was assessed afterward in a separate test; individuals unable to take off were excluded from the analysis. Emitter groups could be reused for up to two consecutive days; overall, five independent groups of mated females, six of virgin females, and seven of virgin males were used. Between each treatment, the wind tunnel was cleaned with 75% ethanol, and all artificial plants were replaced.

To record behavior, we used the event-logging software The Observer (version 10.5; Noldus Information Technology, Wageningen, the Netherlands). For each receiver tested, we recorded whether the wasp took off (binary variable), its takeoff latency (i.e., the time elapsed between reaching the opening of the release tube and takeoff), the duration of flight, and whether the insect reached the odor source (i.e., landed on the plant next to the flask containing emitters). Time variables were measured in seconds with a precision of 0.1 s. In addition, a random subset of 241 individuals from all categories was freeze-killed after testing, and total body length was measured using a digital microscope (Keyence VHX-2000).

### Field experiment

The experiment consisted of releasing *M. ridens* males in an apple orchard and recording their subsequent occurrence on traps baited with virgin females, mated females, or no females. Traps were placed in two apple trees: the focal tree where males were released and a distant tree located 10 m away. The orchard was small (approximately 20 trees), largely unmanaged, and located next to the laboratory (43°36′54″N, 7°04′53″E). Traps consisted of a plastic ring (5 cm diameter, 0.5 cm thick) sandwiched between two layers of thin metal mesh. At the beginning of each three-day trial, traps were baited with five 2– to 3-day-old virgin or mated females and maintained overnight under laboratory conditions with honey and water. Control traps were prepared identically but contained no females. Each morning, three traps of each type were hung in the focal tree and one of each type in the distant tree. Within trees, trap positions were randomized, 1.5–2 m above ground and at least 50 cm apart. At the end of each daily observation period, traps were returned to the laboratory and maintained overnight with honey and water. Female survival was checked daily, and dead females were replaced as needed to maintain five females per trap throughout the experiment.

We visually recorded the number of male *M. ridens* present on each trap every 5 min over three successive days: the day before release, the day of release, and the following day. Observation periods generally extended from approximately 09:00 to 12:00 before release and from approximately 09:00 to 18:00, with a midday interruption, on the two subsequent days. On the day of release, approximately 100 males, all <7 days old, were released at the base of the focal tree trunk. Males observed on traps were not captured or marked, so the same individual could be recorded multiple times. Temperature and relative humidity were recorded at 15-min intervals throughout the observation periods. The experiment was repeated five times between September and October 2022.

### Chemical and electrophysiological analyses

All analyses were conducted at the Platform for Chemical Analyses in Ecology (PACE), a technical facility of the LabEx CeMEB (Centre Méditerranéen pour l’Environnement et la Biodiversité, Montpellier, France). Volatile organic compounds (VOCs) released by *Mastrus ridens* were collected using three complementary methods: solvent extraction, static headspace sampling by solid-phase microextraction (SPME), and dynamic push–pull headspace extraction. Insect category (virgin males [VM], virgin females [VF], and mated females [MF]) and age (0–2, 3–5, and >6 days) varied across experiments. For each sampling batch, a blank control without insects was collected.

#### Solvent extraction

Virgin females were briefly chilled at 4 °C for 5 min and then immediately transferred in groups of five into 3 mL amber vials containing 500 μL of solvent (hexane, dichloromethane, or methanol). Extractions were performed on virgin females from three age classes (0–2, 3–5, and >6 days), with one replicate for each age × solvent combination. In addition, groups of 20 females comprising both virgin and mated individuals, all older than two weeks, were extracted separately with each of the three solvents. After 10 min of extraction, the bodies were removed and the extracts were concentrated under a gentle stream of nitrogen to approximately 200 μL and stored at −20 °C. Dichloromethane and methanol extracts did not reveal additional volatile compounds compared with hexane extracts, and subsequent analyses therefore focused on hexane extracts.

#### Solid-phase microextraction (SPME)

Volatiles were collected from both sexes using solid-phase microextraction (SPME). For each replicate, a group of 20 parasitoids (mated females, virgin females, or virgin males) was introduced into a 10 mL glass vial containing a 1 × 1 cm piece of filter paper moistened with 20 μL of water. The vial was sealed with a silicone plug pierced by a perforated PTFE tube extending to the bottom of the vial, the upper end of which was closed with Nalophan film (Kalle Nalo GmbH, Wursthüllen, Germany). After a 5-min acclimatization period, the Nalophan film was pierced and a 65 μm polydimethylsiloxane/divinylbenzene (PDMS/DVB) SPME fiber (Supelco, Bellefonte, PA, USA) was inserted into the vial and exposed for 2 h. A total of 11 independent replicates were performed with mated females, 10 with virgin females, and 7 with virgin males.

#### Dynamic headspace collection

To maximize the quantity of putative VOCs collected, groups of insects were monitored for several consecutive days. Due to rearing constraints and uncertainty regarding the number of wasps required to detect volatile emissions, two approaches were used: (1) combining all available females, regardless of mating status and age (0–10 days at the start of collection), to form two independent batches of 50 and 100 individuals, each monitored for three consecutive days (resulting in three collections per batch); and (2) assembling two independent batches of 30 freshly emerged virgin females (0–1 and 0–2 days old), which were monitored daily for 5 and 4 days, respectively (resulting in a total of nine daily collections).

Volatiles were collected using 100 mL glass vials sealed with silicone plugs pierced by syringe needles and containing a 2 × 2 cm piece of filter paper moistened with 60 μL of water. Up to three vials were sampled simultaneously, including blank controls containing no insects. Headspace collection followed a method adapted from Proffit et al. (2020), originally developed for plants but here modified for arthropods. Air was pumped into the system at an inlet flow rate of 300 mL·min⁻¹ and removed at 200 mL·min⁻¹, with both flow rates regulated by flowmeters. The higher inlet than outlet flow maintained a slight positive pressure, thereby limiting the entry of potentially contaminated ambient air. Inlet air was purified through a charcoal filter, whereas outlet air passed through Porapak-Q sorbent traps (VCT glass tubes, ARS, Gainesville, Florida, USA) to capture VOCs. Each collection lasted 7 h. Porapak-Q traps were initially eluted sequentially with 150 μL each of hexane, dichloromethane, and methanol. As dichloromethane and methanol did not reveal additional compounds, subsequent samples were eluted with hexane only.

#### Gas chromatography-mass spectrometry (GC-MS)

All samples were analyzed using a Shimadzu QP2010 Plus gas chromatograph–mass spectrometer (GC–MS) equipped with a quadrupole mass analyzer (Shimadzu Scientific Instruments, Kyoto, Japan). The GC was fitted with an Optima 5-MS fused-silica capillary column (30 m × 0.25 mm internal diameter, 0.25 μm film thickness; Macherey-Nagel, Düren, Germany), with helium as the carrier gas at a constant flow rate of 1 mL·min⁻¹. SPME fibers were thermally desorbed directly in the injector at 250 °C in spitless mode, whereas 1 μL of liquid extracts or eluates was injected for GC–MS analysis. The oven temperature was programmed from 40 °C (1-min hold) to 220 °C at 12 °C·min⁻¹, followed by a final 2-min hold. Data were processed using GCMSsolution software (Shimadzu Scientific Instruments, Kyoto, Japan).

Retention times of a standard mixture of n-alkanes (Alkanes standard solution 04070, Sigma-Aldrich) were used to calculate linear retention indices (LRIs). Chromatograms obtained from parasitoid samples were systematically compared with their corresponding blank controls, and peaks also present in the controls were excluded from consideration as parasitoid-associated compounds.

Compound identification was based on comparison of mass spectra with the NIST 2007 and 2011 libraries and the Wiley 9th edition library, comparison of retention times and mass spectra with external synthetic standards when available, and comparison of LRIs with published values (Adams 2007).

#### Gas chromatography–electroantennographic detection (GC–EAD)

Electroantennographic recordings were made on adult *Mastrus ridens* males using coupled gas chromatography–electroantennographic detection (GC–EAD). Each male was immobilized, and its head was severed and inserted into a customized glass capillary. The capillary (76 mm long, 1.12 mm diameter; World Precision Instruments, Sarasota, USA) was pulled and cut using a vertical micropipette puller (model P-30, World Precision Instruments, Hertfordshire, UK). It was filled with Ringer’s solution (6 g·L⁻¹ NaCl, 0.4 g·L⁻¹ KCl, 0.3 g·L⁻¹ CaCl₂, and 3 g·L⁻¹ NaHCO₃) and connected to the reference electrode. The distal tip of one antenna was placed in contact with the recording electrode, which was filled with the same electrolyte.

A stimulus controller (CS-55, Syntech, Ockenfels, Kirchzarten, Germany) delivered a continuous flow of purified and humidified air at 245 mL·min⁻¹ through a glass tube carrying the GC effluent to the antenna. A 2 μL sample was injected into an Agilent 8890 gas chromatograph (Agilent, Santa Clara, USA) equipped with a flame ionization detector (FID) and a split/splitless multimode inlet maintained at 200 °C (split ratio 1:20). Separation was achieved using an Optima 5-MS capillary column (30 m × 0.25 mm internal diameter, 0.25 μm film thickness; Macherey-Nagel, Düren, Germany), with helium as the carrier gas at 1 mL·min⁻¹. The oven temperature was programmed from 40 °C (1-min hold) to 220 °C at 12 °C·min⁻¹, followed by a final 2-min hold.

The column effluent was split between two deactivated fused-silica capillaries (100 cm × 0.25 mm), with 10% directed to the FID maintained at 300 °C and 90% routed through a heated EAD transfer line maintained at 200 °C (Syntech). Antennal responses were digitized using an IDAC-2 acquisition controller (Syntech) and processed using GcEad 1.2.5 software (Syntech) (Proffit et al. 2020). Seventeen males were tested in total, three of them twice depending on signal quality, yielding 12 recordings with pooled Porapak-Q extracts and eight with solvent extracts. EAD-active peaks were identified from measurable antennal depolarization responses and subsequently matched to corresponding peaks characterized by GC–MS using retention indices and mass spectra.

### Statistical analyses

Statistical analyses were conducted in the R environment (version 4.5.1, Great Square Root; R Core Team 2025). All central tendencies reported in the Results section correspond to model-estimated marginal means, accompanied by 95% confidence intervals. Annotated R Markdown files are provided as supplementary materials.

#### Wind tunnel experiment

We fitted generalized linear models (GLMs) and survival models to analyze four behavioral variables: takeoff probability, takeoff latency, flight duration, and the probability of reaching the odor source. Takeoff probability and the probability of reaching the odor source were modeled using binomial GLMs with a logit link. Bias-reduced estimation (brglmFit option) was used to handle complete separation caused by some factor combinations in which all tested individuals took off, resulting in no within-group variance. Takeoff latency was analyzed using a Weibull survival model, treating individuals that did not initiate flight within the 5-minute observation window as right-censored. Flight duration was analyzed using a Gamma GLM with a log link, restricted to individuals that took off.

For each behavioral variable, we fitted a series of nested models to the full dataset to test for effects of receiver type, emitter type, and their interaction. Main effects were tested by comparing single-factor models to the null (intercept-only) model, while the interaction was tested by comparing the full interaction model to the additive model. Likelihood ratio tests (LRT) were used in all cases, assuming a χ² distribution of the test statistic. When a significant interaction was detected, separate models were fitted for each receiver type to clarify emitter effects within each subset. Post-hoc comparisons were conducted using Bonferroni-adjusted pairwise tests. Although conservative, this correction was preferred for its robustness in controlling type I error.

#### Field data

Raw field data consisted of 13,980 visual counts of M. ridens males recorded every 5 min on 12 traps (3 trap types × 4 traps per type) over three consecutive days in each of five replicated experiments. Because males were not marked and could therefore be recorded repeatedly at successive censuses, and because raw counts contained a large excess of zeros, observations were aggregated into hourly presence/absence for each trap. Presence was scored as 1 when at least one male was observed during the corresponding hourly period and 0 otherwise. Pre-release observations were used as baseline information but were excluded from the inferential analysis, which therefore included the day of release and the following day. Each trap type (VF, MF, CT) was represented by three traps on the focal tree and one trap on a distant tree located 10 m away.

Hourly presence/absence data were analyzed using generalized linear mixed models (GLMMs) with a binomial error distribution and logit link. The full model included trap type (VF, MF, CT), trap distance (near vs. far), day (day of release vs. following day), and hourly slot (1–8) as fixed effects. Replicate experiment and individual trap within replicate experiment were included as random intercepts to account for repeated observations of the same traps. Candidate models comprising all combinations of these main effects were ranked according to AICc, and the model with the lowest AICc was retained. The contribution of each retained fixed effect was then assessed by likelihood ratio tests, comparing the selected model with the corresponding model lacking that effect.

#### Chemical analyses

Differences in the occurrence of each compound among extraction methods, female mating-status categories, and sexes were assessed using Fisher’s exact tests. Comparisons of headspace sampling methods reported in the Results were restricted to SPME and dynamic headspace samples. Comparisons between sexes were restricted to SPME samples, the only sampling method providing replicated samples for both males and females.

## Results

### Orientation toward conspecifics in the wind tunnel

After exiting the release tube and thereby being exposed to conspecific volatiles, wasps tested as receivers could take off and fly in the wind tunnel. Among the 339 individuals tested, three-quarters (0.75 [0.71, 0.80]) took off within the five minutes of post-release observation. Takeoff probability was strongly affected by receiver type (LRT = 33.1, df = 2, p < 0.0001; Fig. 2). Regardless of putative volatiles, males took off more frequently (0.92 [0.86, 0.96]) than virgin and mated females (0.70 [0.61, 0.78] and 0.63 [0.54, 0.71], respectively). The type of emitter had no significant main effect (LRT = 4.75, df = 3, p = 0.19), and the interaction between receiver and emitter types was only marginally significant (LRT = 12.6, df = 6, p = 0.051). There was a tendency for males to take off more often when exposed to odors from virgin or mated females, whereas females took off independently of emitter identity (Fig. 2). Consistently, *all* males exposed to virgin or mated females did take flight, a striking result observed in no other receiver–emitter combination.

**Figure 2.**
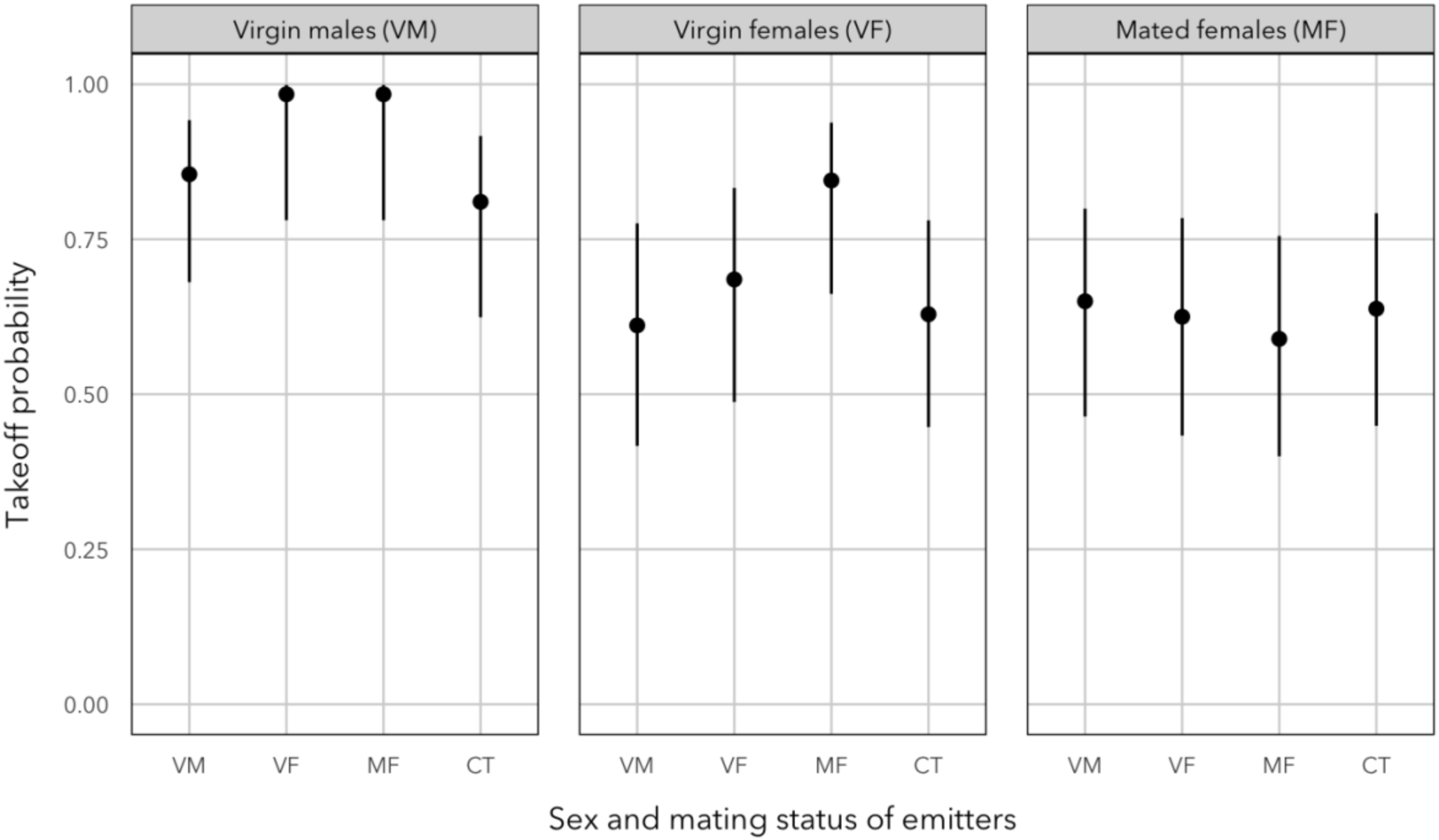
Effect of sex and mating status of individual *Mastrus ridens* used as emitters on the probability that conspecific receivers take off. Emitter types include virgin males (VM), virgin females (VF), mated females (MF), and a control with no emitter (CT). Each panel corresponds to a receiver type: virgin male, virgin female, or mated female. Points represent estimated marginal probabilities (± 95% confidence intervals) obtained from a full binomial generalized linear model (receiver × emitter) fitted to the whole dataset.

Takeoff latency varied markedly depending on the combination of receiver and emitter types (interaction: LRT = 26.6, df = 6, p < 0.0001; Fig. 3). Among males, emitter identity had a strong effect (LRT = 33.8, df = 3, p < 0.0001): males took off much more rapidly when exposed to virgin or mated females (6 s [3, 12] and 8 s [4, 16], respectively) than when exposed to other males or to the control condition (60 s [30, 119] and 70 s [33, 145], respectively). In contrast, takeoff latency in virgin and mated females was unaffected by emitter identity (VF: LRT = 1.89, df = 3, p = 0.60; MF: LRT = 1.67, df = 3, p = 0.64). Regardless of emitter identity and mating status, female latency averaged 234 s [177, 309], which was substantially longer than that of males (main effect of receiver: LRT = 76.3, df = 2, p < 0.0001).

**Figure 3.**
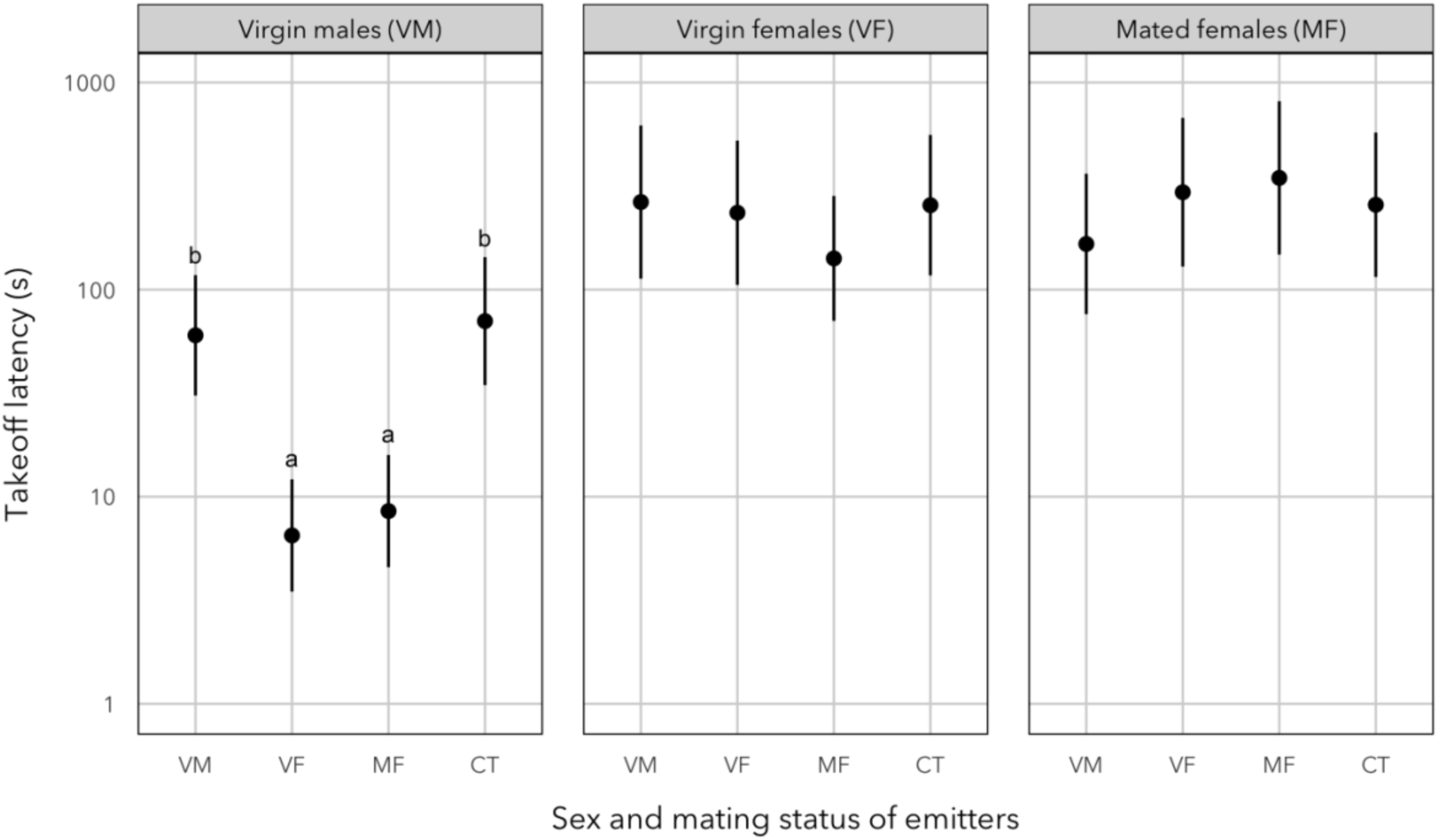
Effect of sex and mating status of individual *Mastrus ridens* used as emitters on receiver takeoff latency. Emitter types include virgin males (VM), virgin females (VF), mated females (MF), and a control with no emitter (CT). Each panel corresponds to a receiver type: virgin male, virgin female, or mated female. Points represent estimated marginal means (± 95% confidence intervals) obtained from a full Weibull survival model (receiver × emitter) fitted to the whole dataset. Estimates incorporate censored observations corresponding to wasps that did not take off within the 300 – second post – release observation period. The y–axis is on a logarithmic scale. Letters indicate statistically distinct groups based on Bonferroni–adjusted pairwise comparisons (α = 0.05), performed only for male receivers in which the emitter effect was significant.

For the 256 individuals that took off, flight duration was highly variable. The duration of the first flight ranged from less than one second to nearly 200 seconds, with a median of 5.3 s. As for takeoff latency, the interaction between receiver and emitter types was significant (LRT = 33.3, df = 6, p < 0.0001), indicating that the locomotor response to conspecific volatiles depended on the sex and mating status of the receivers. Among males, emitter identity had a strong effect (LRT = 27.6, df = 3, p < 0.0001): they flew longer when exposed to odors from virgin or mated females (26 s [17, 40] and 22 s [14, 34]) than when exposed to other males or to an absence of conspecifics (11 s [7, 17] and 7 s [4, 12]; Fig. 4). In contrast, emitter identity had no effect on female flight duration (VF: LRT = 5.70, df = 3, p = 0.13; MF: LRT = 6.51, df = 3, p = 0.09). Females flew an average of 5.2 s [4.6, 6.0] regardless of their mating status and volatiles from conspecifics (Fig. 4).

**Figure 4.**
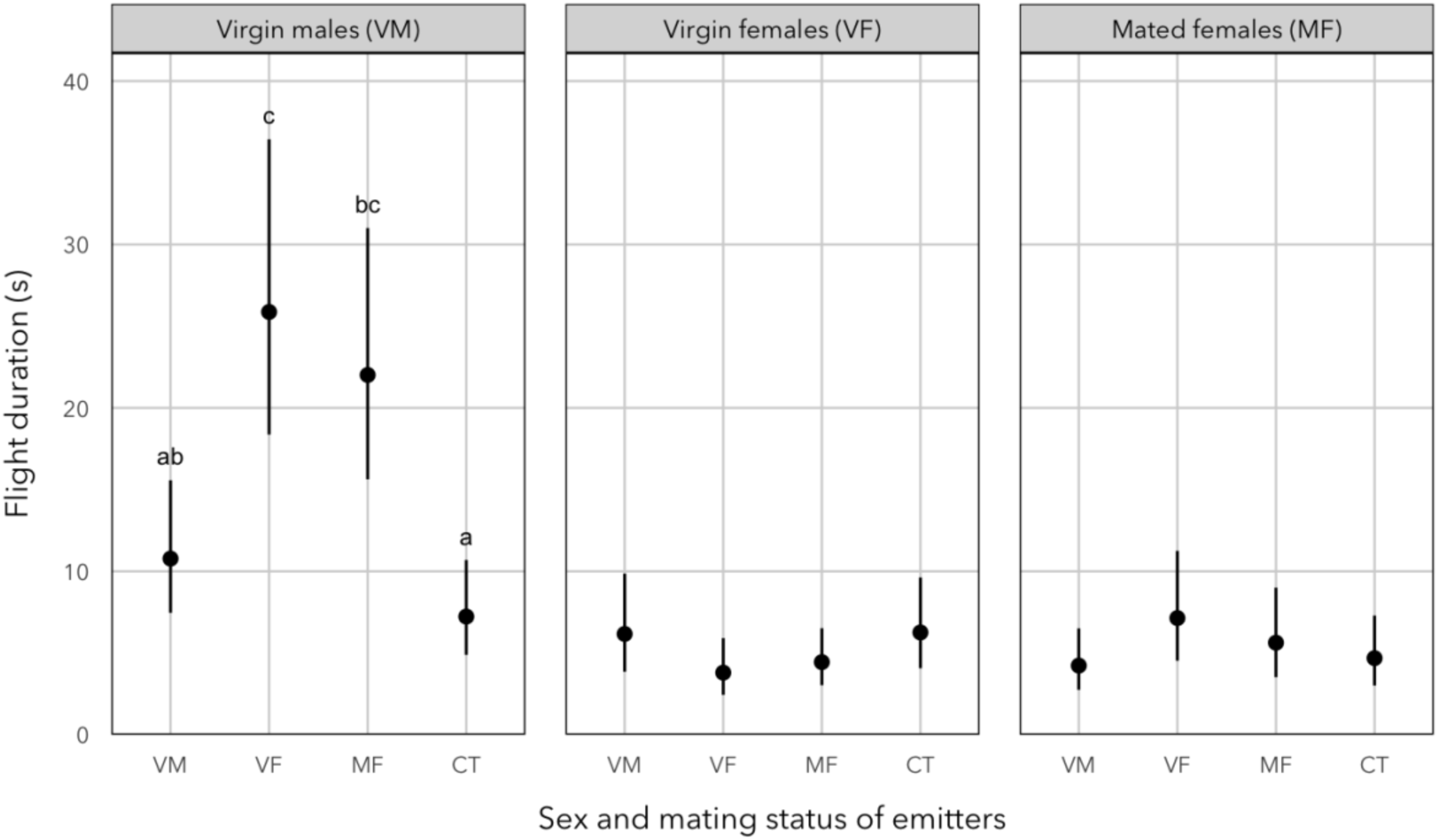
Effect of sex and mating status of individual *Mastrus ridens* used as emitters on flight duration in receivers. Emitter types include virgin males (VM), virgin females (VF), mated females (MF), and a control with no emitter (CT). Each panel corresponds to a receiver type: virgin male, virgin female, or mated female. Points represent estimated marginal means (± 95% confidence intervals) obtained from a full Gamma generalized linear model (receiver × emitter) fitted to the whole dataset. Letters indicate statistically distinct groups based on Bonferroni – adjusted pairwise comparisons (α = 0.05), performed only for male receivers in which the emitter effect was significant.

Reaching the source of volatiles represents the ultimate criterion of successful orientation. Among the 339 wasps tested, 46 reached the odor source (13.6%) but as with some other behavioral variables, the interaction between receiver and emitter types was significant (LRT = 20.6, df = 6, p = 0.002). Emitter identity strongly influenced the probability of reaching the source in males (LRT = 56.6, df = 3, p < 0.0001), but had no effect in females, whether virgin or mated (VF: LRT = 0.17, df = 3, p = 0.98; MF: LRT = 2.50, df = 3, p = 0.48). As illustrated in Fig. 5, males reached the emitters much more frequently when exposed to virgin or mated females (0.60 [0.47, 0.71]) than when exposed to other males or to control flasks (0.04 [0.01, 0.14]). In contrast, the probability of reaching the source in females was remarkably low (0.04 [0.02, 0.07]) and unaffected by the olfactory context.

**Figure 5.**
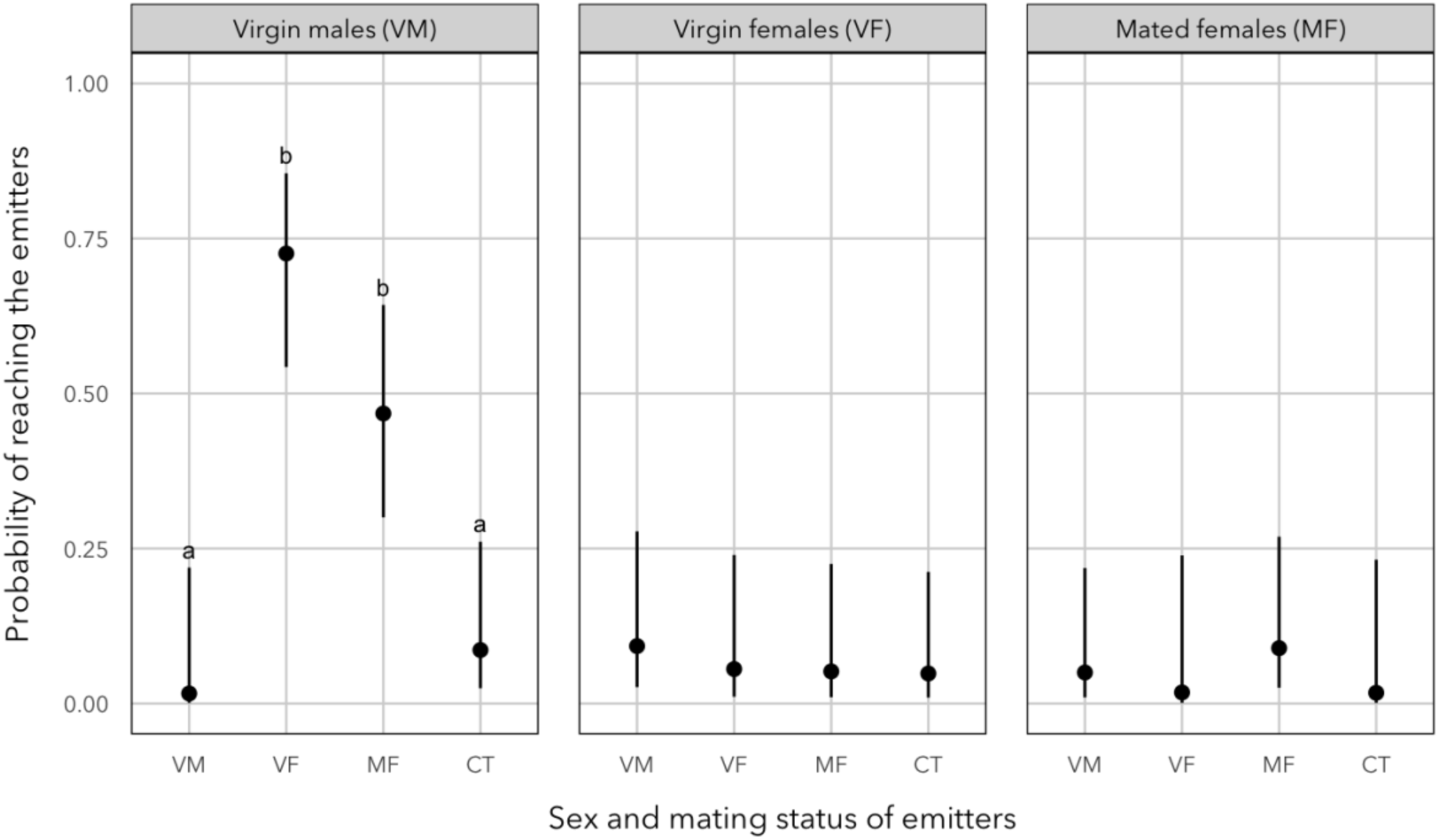
Effect of sex and mating status of *Mastrus ridens* individuals used as emitters on the probability that conspecific receivers reach the emitters. Emitter types include virgin males (VM), virgin females (VF), mated females (MF), and a control with no emitter (CT). Each panel corresponds to a receiver type: virgin male, virgin female, or mated female. Points represent estimated marginal probabilities (± 95% confidence intervals) obtained from a full binomial generalized linear model (receiver × emitter) fitted to the whole dataset. Letters indicate statistically distinct groups based on Bonferroni – adjusted pairwise comparisons (α = 0.05), performed only for male receivers in which the emitter effect was significant.

Altogether, these results suggest that males respond to volatiles emitted by females, whether virgin or mated: when downwind from females, they are more likely to take off, they do so sooner, they fly longer, and they are more likely to land on the plant next to the source of volatiles. We therefore restricted the analysis of body size to males exposed to females. Both whole body length and antenna length were negatively associated with takeoff latency, indicating that larger males initiated flight sooner (body: LRT = 7.93, df = 1, p = 0.0048; antenna: LRT = 6.58, df = 1, p = 0.010). In contrast, neither measure of size influenced flight duration or the probability of reaching the source.

### Male orientation toward females in the field

After release in the apple orchard, males were frequently observed approaching and hovering in front of female-baited traps (Fig. 6). Model selection strongly favored the full binomial GLMM including trap type, distance, day, and time slot (AICc = 686.3, Akaike weight = 0.893). The next-best model, which excluded distance, had substantially less support (ΔAICc = 4.30, weight = 0.104). Likelihood ratio tests confirmed strong effects of trap type (χ² = 107.4, df = 2, p < 0.0001), day (χ² = 13.7, df = 1, p < 0.001), time slot (χ² = 18.1, df = 1, p < 0.0001), and a weaker but significant effect of distance (χ² = 6.33, df = 1, p = 0.012).

**Figure 6.**
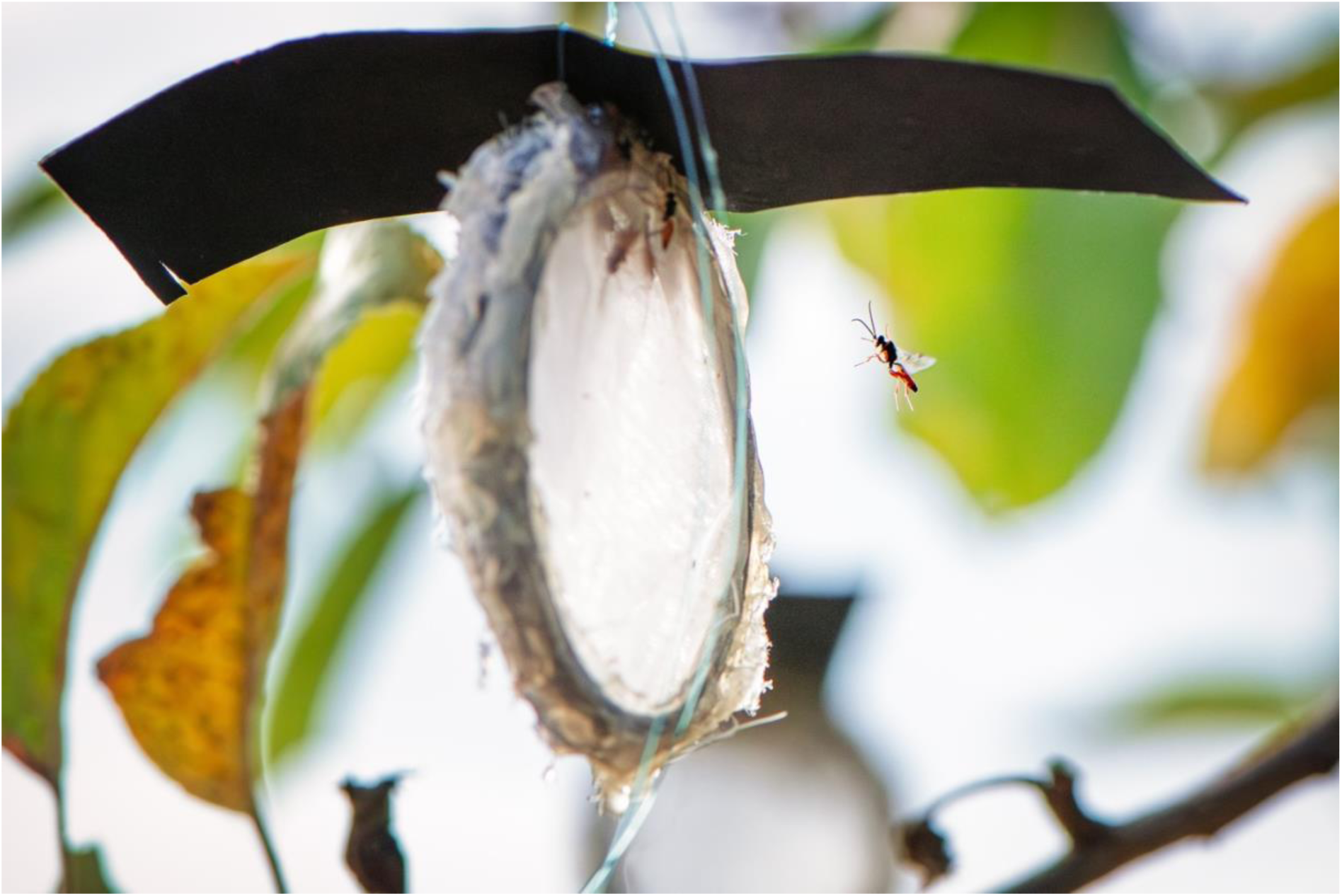
Male *Mastrus ridens* in flight approaching a female-baited trap in an apple orchard during the field experiment. The male is shown hovering near the trap with antennae extended. Traps consisted of a plastic ring (5 cm diameter) sandwiched between two layers of metal mesh and were suspended 1.5–2 m above ground. Approximately 100 males were released at the base of a focal tree, and male presence on traps was visually recorded at 5-min intervals over three consecutive days under natural field conditions.

Male presence differed sharply among trap types (FDR-adjusted pairwise tests, all p < 0.0001). The probability of observing at least one male was highest on virgin-female traps (0.88 [0.65–0.96]), intermediate on mated-female traps (0.41 [0.16–0.72]), and very low on controls (0.015 [0.004– 0.061]). The contrast between virgin and mated females was striking: males were more than twice as likely to be observed on virgin-female traps as on mated-female traps (Fig. 7), a difference far stronger than in wind-tunnel assays.

**Figure 7.**
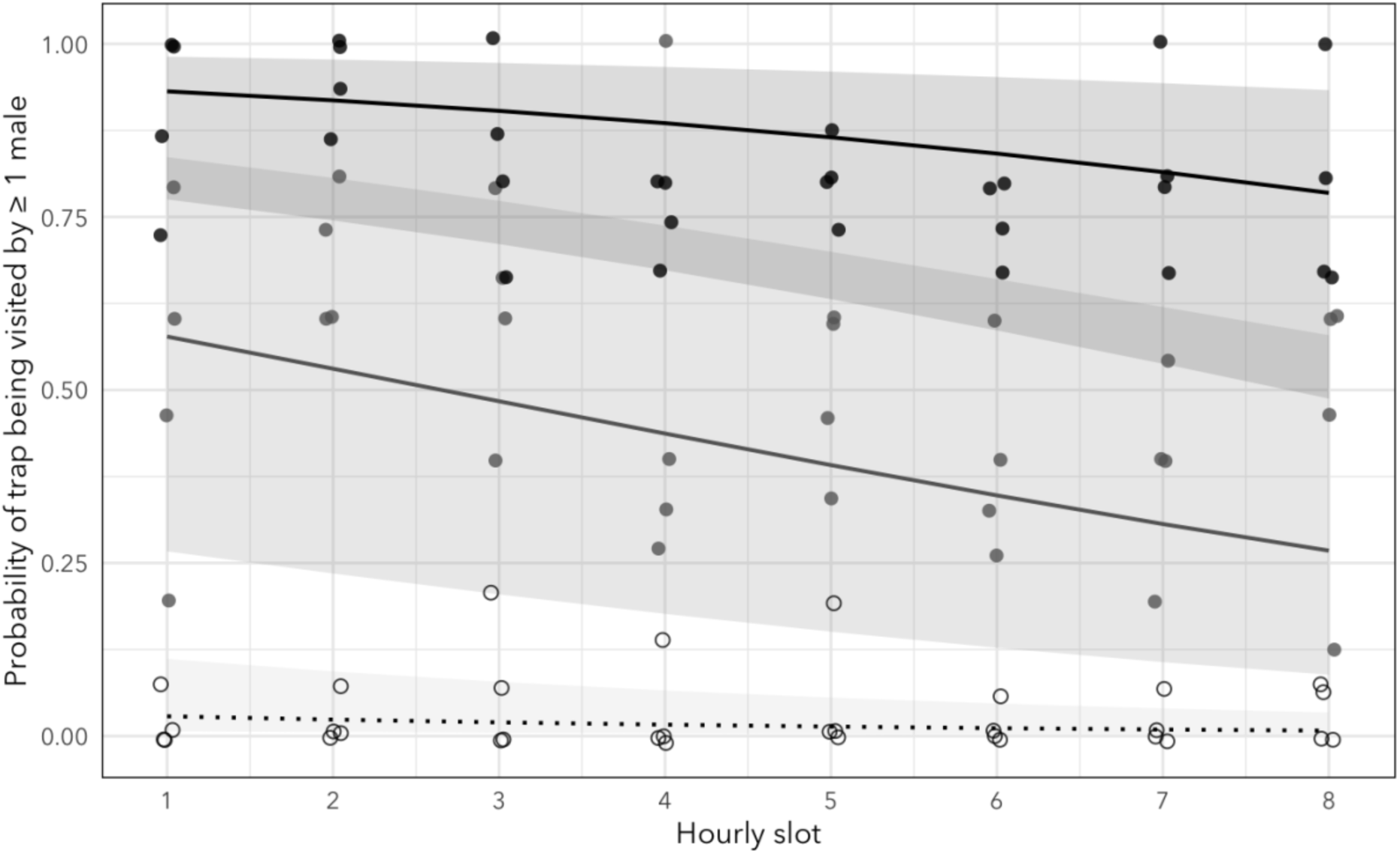
Observed and model-estimated hourly probabilities of male Mastrus ridens presence on traps. Black filled circles and the solid black line represent traps baited with virgin females; grey filled circles and the solid grey line represent traps baited with mated females; open circles and the dotted black line represent empty control traps. For each trap and hourly slot, presence was scored as 1 if at least one male was observed during any of the 12 successive 5-min censuses, and as 0 otherwise. Points show observed proportions calculated separately for each combination of day relative to male release (release day or following day), distance from the release point (near or far), trap type, and hourly slot. Points are slightly jittered for visibility. Lines show predicted probabilities, and shaded bands indicate 95% confidence intervals, from the selected binomial generalized linear mixed model. The model included trap type, day, distance, and hourly slot as fixed effects, with random intercepts for replicate experiment and individual trap. Predictions are marginalized over day and distance.

Male presence decreased over the course of the day (Fig. 7) and was lower on the day following release than on the release day. Unexpectedly, male presence was higher on distant traps, located 10 m from the release tree, than on traps placed on the focal tree (0.45 [0.17–0.76] vs. 0.26 [0.09–0.55], respectively).

### Identification of the volatile compounds emitted

Because our study focused on volatile compounds potentially involved in attraction from a distance, we restricted our analyses to biogenic volatile organic compounds (VOCs) with molecular masses between 38 and 350 amu. Across all extraction methods combined (solvent extraction, solid-phase microextraction [SPME], and dynamic headspace), a total of 40 female samples were analyzed. Comparative GC–MS analyses consistently revealed three target compounds: 4-methyl-3-hexanol, 4-methyl-3-heptanone, and methyl 6-methylsalicylate (Table 1). Among these, methyl 6-methylsalicylate was the most frequently detected, occurring in 33 samples (83% [67–93]) across all extraction methods. In contrast, 4-methyl-3-heptanone was detected in 22 samples (55% [38–71]) and 4-methyl-3-hexanol in 14 samples (35% [21–52]).

**Table 1.**
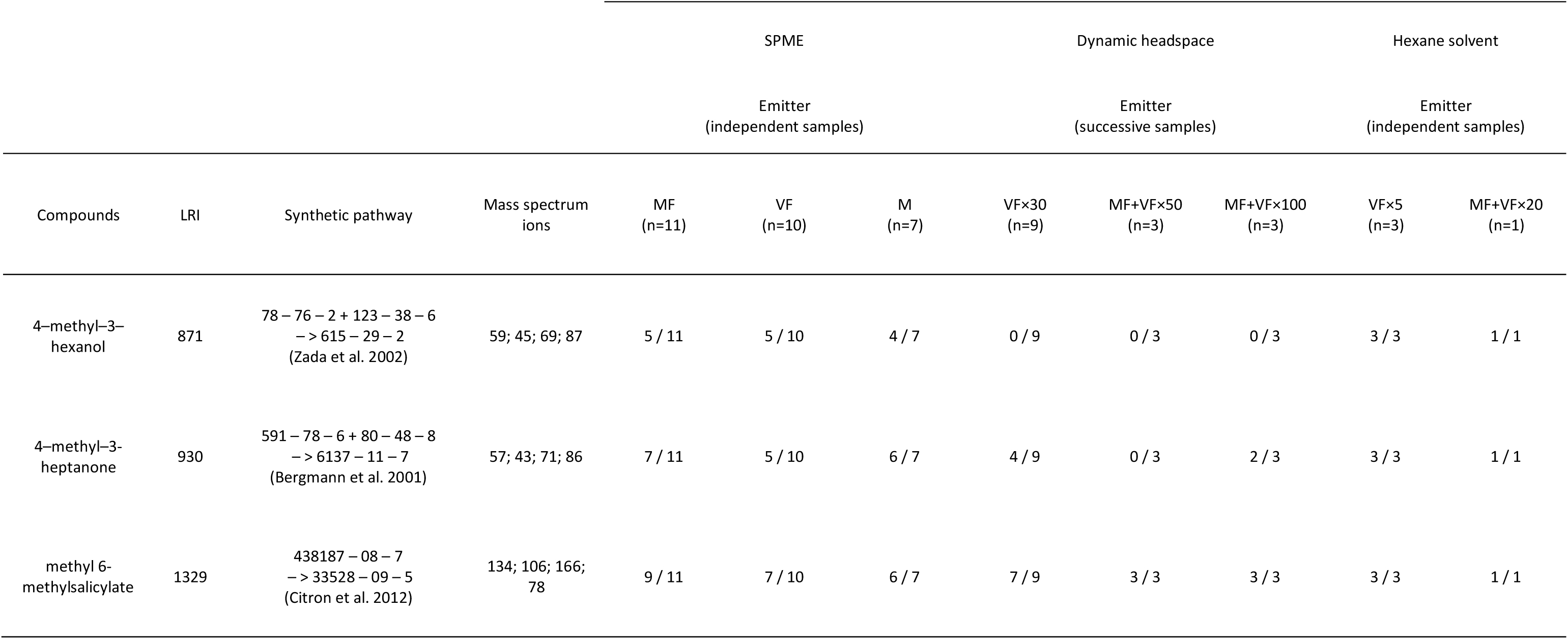
Volatile organic compounds (VOCs) identified by GC–MS in *Mastrus ridens* samples collected using three complementary methods: solid-phase microextraction (SPME), dynamic headspace extraction (Porapak-Q), and solvent extraction with hexane. For each compound, the table reports its linear retention index (LRI), diagnostic mass spectrum ions (m/z), and a representative synthetic pathway (cited by CAS numbers and references). The number of samples with detection over the number of samples analyzed is shown for each compound across emitter groups and sampling methods. Emitter types include virgin females (VF), mated females (MF), mixed groups containing both virgin and mated females (VF + MF), and males (M). Sample sizes (n) indicate the number of independent or successive replicates per group. VOCs were considered present when detected above background in comparison with control samples. References listed for each compound correspond to prior chemical or biosynthetic studies that report the occurrence or synthesis of these molecules in natural systems. References cited in Table 1 Bergmann, J., C. Löfstedt, Vladimir D. Ivanov, and W. Francke. 2001. Identification and Assignment of the Absolute Configuration of Biologically Active Methyl-Branched Ketones from Limnephilid Caddis Flies. European Journal of Organic Chemistry **2001**:3175–3179. Citron, C. A., P. Rabe, and J. S. Dickschat. 2012. The scent of bacteria: headspace analysis for the discovery of natural products. J Nat Prod **75**:1765–1776. Zada, A., V. Soroker, M. Harel, J. Nakache, and E. Dunkelblum. 2002. Quantitative GC analysis of secondary alcohol pheromones: determination of release rate of red palm weevil, Rhynchophorus ferrugineus, pheromone from lures. J Chem Ecol **28**:2299–2306.

All three compounds were consistently detected in hexane extracts (4/4). Comparisons restricted to SPME and dynamic headspace samples revealed that 4-methyl-3-hexanol was detected significantly more frequently in SPME samples (10/21) than in dynamic headspace samples (0/15; Fisher’s exact test, p = 0.0018). In contrast, occurrence did not differ between the two sampling methods for 4-methyl-3-heptanone (12/21 vs. 6/15; p = 0.50) or methyl 6-methylsalicylate (16/21 vs. 13/15; p = 0.67).

No effect of female mating status was detected on the occurrence of any of the three compounds (Fisher’s exact tests, all p > 0.45). We detected 4-methyl-3-hexanol in 8/22 virgin, 5/11 mated, and 1/7 mixed-status female extracts. For 4-methyl-3-heptanone, the corresponding frequencies were 12/22, 7/11, and 3/7, respectively. Methyl 6-methylsalicylate remained the most consistently detected compound, occurring in 17/22 virgin, 9/11 mated, and 7/7 mixed-status female extracts.

Because only SPME provided replicated samples for both sexes, comparisons between males and females were restricted to this sampling method. No effect of sex was detected on the occurrence of any of the three compounds (Fisher’s exact tests, all p ≥ 0.36). We detected 4-methyl-3-hexanol in 10/21 female and 4/7 male extracts, and 4-methyl-3-heptanone in 12/21 female and 6/7 male extracts. Methyl 6-methylsalicylate was again the most consistently detected compound, occurring in 16/21 female and 6/7 male extracts.

Some SPME chromatograms also contained several n-alkanes, most likely corresponding to slightly volatile cuticular hydrocarbons. These compounds were not considered further as candidates for distance-mediated attraction, because their occurrence may have resulted from occasional physical contact between the insects and the SPME sampling device despite the protective PTFE tube.

Analyses of dichloromethane and methanol extracts yielded no additional compounds beyond those already detected in hexane. Across the three solvents, methyl 6-methylsalicylate remained the most consistently detected compound.

### Electroantennographic responses to volatile compounds

GC–EAD recordings from male antennae demonstrated antennal detection of several compounds present in female extracts. A representative coupled GC–EAD trace is shown in Fig. 8, with a clear antennal depolarization coinciding with the FID peak corresponding to methyl 6-methylsalicylate. In pooled Porapak-Q extracts of 0–6-day-old virgin females, methyl 6-methylsalicylate elicited repeatable depolarizations in eight of 12 recordings. In hexane extracts, responses to methyl 6-methylsalicylate were observed in all four recordings with extracts of virgin females and in seven of eight recordings with extracts of mixed batches containing both virgin and mated females.

**Figure 8.**
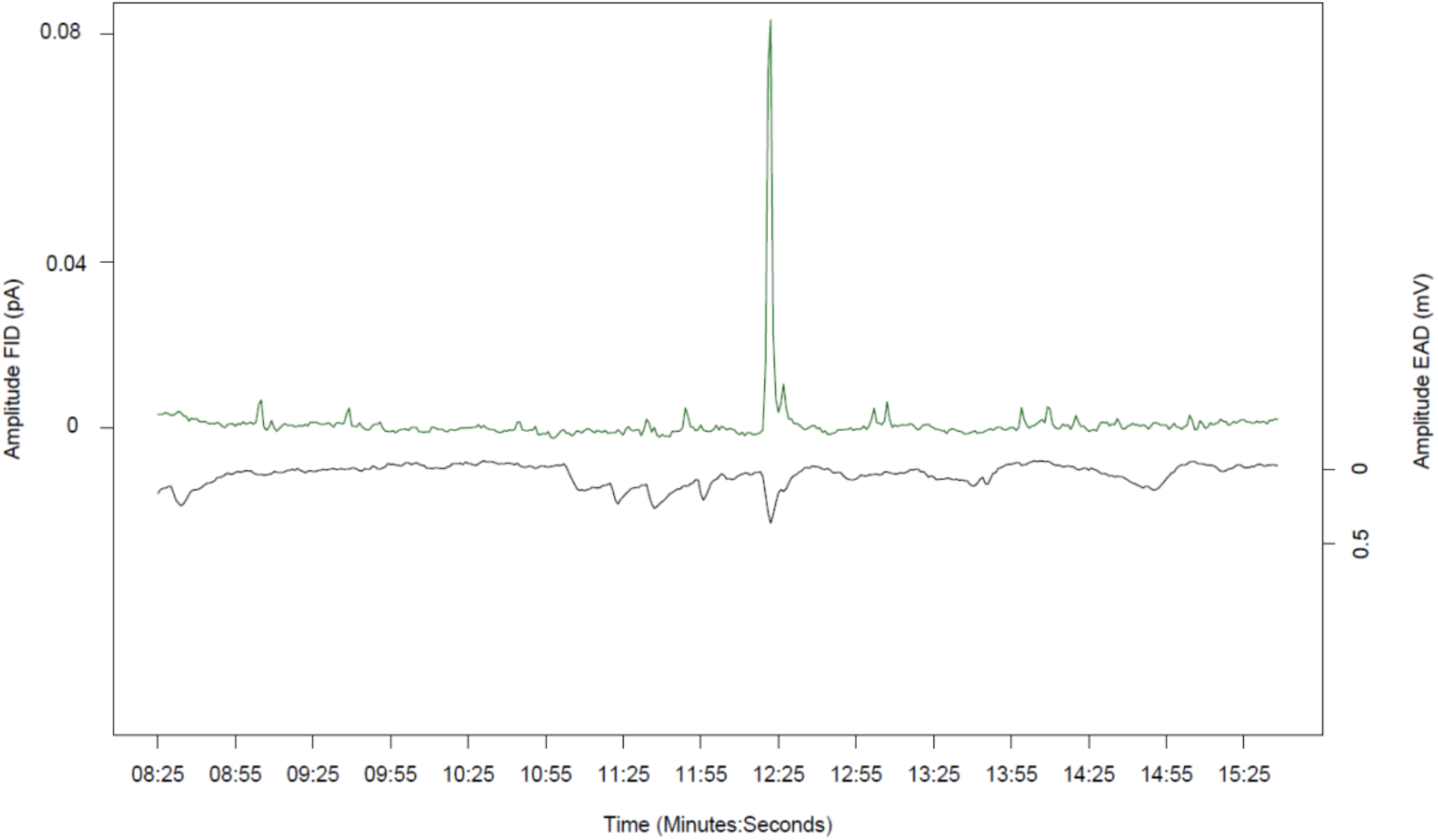
Representative coupled gas chromatography– electroantennographic detection (GC–EAD) trace showing male antennal responses to female headspace volatiles. The upper trace (green) shows the flame ionization detector (FID) signal of a hexane extract from females of mixed mating status. The lower trace (black) shows the electroantennographic response of a male *Mastrus ridens* antenna. A clear antennal depolarization was observed at the retention time corresponding to methyl 6-methylsalicylate, a response consistently detected in eight independent recordings.

In mixed-status female extracts, 4-methyl-3-heptanone also elicited weaker antennal responses in three males. In addition, an unidentified compound (linear retention index [LRI] = 1340; major ions: m/z 148, 134, 57, 43, 85) elicited at least three clear antennal depolarizations. This compound was detected in solvent extracts of mixed-status females but not in SPME chromatograms. Its identity remains to be elucidated.

## Discussion

### Sexual communication in *Mastrus ridens*

The wind-tunnel experiment provides the clearest evidence that sexual communication in *M. ridens* is mediated by volatile cues and is strongly sex-specific at the behavioral level. Only males responded to conspecific odor sources, and their responses were exclusively directed toward females. Males consistently exhibited increased responsiveness to female odors, culminating in directed flight and landing near the source — a pattern never observed in response to male or control stimuli. Field observations corroborated this behavioral pattern under natural conditions, with male presence being highest on traps baited with virgin females, intermediate on those baited with mated females, and very low on controls.

The stronger attraction to virgin than to mated females, together with the absence of female responsiveness and the lack of male attraction to other males, supports the interpretation of a female-emitted volatile sex pheromone rather than a general aggregation cue. Chemical and electrophysiological analyses provide complementary evidence for this inference. Across three sampling methods — solvent extraction, solid-phase microextraction, and dynamic headspace collection — methyl 6-methylsalicylate was the most consistently detected candidate compound in female samples and elicited repeatable antennal depolarizations in males.

Taken together, the behavioral evidence highlights a sex-specific volatile communication system in which females attract males from a distance. In the laboratory, males exposed to female odors took off and flew approximately 0.5 m toward the odor source; in the field, some males reached female-baited traps located 10 m from the release point. Here, “attraction from a distance” refers to orientation before physical contact, rather than to a measured maximum detection range.

The chemical and electrophysiological results further identify methyl 6-methylsalicylate as a biologically active candidate component of the signal, but do not establish that this compound alone accounts for attraction or that it is specific to females. These findings are therefore consistent with a volatile sex pheromone whose behavioral activity may depend on a more complex pheromonal blend, as developed below.

### Methyl 6-methylsalicylate and the case for a multicomponent pheromonal bouquet

Methyl 6-methylsalicylate (M6MS) was the most consistently detected compound in *Mastrus ridens* females and elicited the most consistent antennal responses in males. This aromatic ester (C₉H₁₀O₃) is structurally related to methyl salicylate, a widespread plant volatile released in response to herbivory and other stresses (Ozawa et al. 2000). Together, its recurrent detection across extraction methods and its electrophysiological activity make M6MS our strongest chemical candidate. However, we argue that M6MS is best viewed not as the pheromone itself, but as one component of a multicomponent sex pheromone.

Several observations from our study suggest that M6MS alone is not sufficient to trigger male attraction. First, M6MS was also detected in male odor profiles, yet male-emitted volatiles elicited no behavioral response in the wind tunnel. Thus, naturally occurring M6MS was not sufficient to make an odor source attractive. Second, exploratory wind-tunnel trials with synthetic M6MS alone did not elicit obvious male attraction. Although these assays were not designed for formal inference, they provide no evidence that synthetic M6MS is attractive on its own under our experimental conditions. Together, these independent observations support the view that the presence of M6MS alone does not determine attractiveness. Nonetheless, as we did not measure emission rates, we cannot exclude that males and females release different amounts and that such quantitative differences contribute to their contrasting attractiveness. This possibility is biologically plausible: in the parasitoid wasp *Spalangia endius*, previously mated females release more M6MS than virgin females during active courtship, and higher M6MS emissions are associated with reduced male courtship (Mowles et al. 2013). In addition to this possible quantitative effect, our chemical analyses in *M. ridens* revealed other compounds that could contribute to the signal. Two aliphatic compounds, 4-methyl-3-hexanol and 4-methyl-3-heptanone, were detected in both sexes, and the latter elicited antennal responses in a subset of males. In addition, an unidentified compound at LRI 1340 was also EAD-active. These compounds are therefore candidates for the additional chemical information that, together with M6MS, could make female odor attractive.

If M6MS alone is insufficient for attraction, its behavioral effect may depend on the blend in which it occurs. A particularly relevant precedent is provided by the parasitoid wasp *Asobara tabida*, in which M6MS is part of a chemically diverse female sex pheromone. M6MS alone elicits only weak male courtship responses, whereas its combination with other female-derived compounds produces responses comparable to those elicited by female extracts (Stokl et al. 2014). A similarly striking example occurs in *Polyergus ants*, in which M6MS itself is part of a multicomponent sex pheromone. In P. breviceps, M6MS occurs in queens, workers, and males, although at much higher levels in queens, yet M6MS alone is completely unattractive to males; strong attraction requires its combination with 3-ethyl-4-methylpentanol (Greenberg et al. 2004). In *P. rufescens*, M6MS accounts for about 92% of the queen mandibular-gland blend, whereas 3-ethyl-4-methylpentanol represents only about 2%, but strong male attraction likewise requires both compounds (Castracani et al. 2008). Together, these studies show that M6MS can be an important, even quantitatively dominant, pheromone component without being sufficient on its own to elicit the complete behavioral response.

The relevance of blend-dependent orientation is also supported, more indirectly, by the olfactory ecology of *M. ridens* itself. Females locate aggregated *Cydia pomonella* larvae by exploiting the host aggregation pheromone as a long-range kairomone (Jumean et al. 2009). A synthetic 11-component blend reproduced attraction, and deletion experiments showed that only three components could be removed without loss of attractiveness, leaving an eight-component active blend (Jumean et al. 2005). Thus, orientation in *M. ridens* can depend on information distributed across multiple odor components. Although this host-location system is chemically and functionally distinct from sexual communication, it provides a within-species precedent for blend-dependent orientation.

These specific examples fit a broader pattern in parasitoid sexual communication, in which behavioral information can reside in the composition and relative proportions of a pheromonal blend rather than in the presence of a single major compound. In *Leptopilina heterotoma*, for example, the major pheromone component (-)-iridomyrmecin is ineffective alone; reliable male attraction requires several minor iridoid components, and closely related species produce the same compounds in different ratios (Weiss et al. 2013). More generally, combinations of major and minor pheromone components and their relative proportions can contribute to mate recognition in parasitoids (Tonga et al. 2026). The importance of such blend coding has also been demonstrated experimentally in the highly specialized *Ficus carica*–*Blastophaga psenes* interaction, where attraction depends on particular proportions of otherwise common volatile compounds (Proffit et al. 2020).

More broadly, these examples caution against assigning a fixed behavioral meaning to individual semiochemicals. M6MS itself illustrates this point: it contributes to alarm and trail signaling in ants (Duffield and Blum 1975, Morgan and Ollett 1987) and to courtship modulation in parasitoid wasps (Nichols et al. 2010, Stokl et al. 2014). This interspecific functional diversity exemplifies semiochemical parsimony, whereby the same molecule can be coopted into distinct communicative roles (Wyatt 2014).

Taken together, our results argue against M6MS acting alone as the male attractant in *M. ridens*. Instead, they support a model in which M6MS is a key component of a multicomponent pheromonal signal whose behavioral effect depends on its chemical context and may also vary with emission rate. Antennal responses to 4-methyl-3-heptanone in a subset of males, together with the additional unidentified EAD-active compound at LRI 1340, identify plausible additional elements of this signal. The critical next step is therefore to reconstruct candidate blends and test their behavioral activity across controlled component ratios and release rates.

### Sex pheromones, dispersal and the mating system

Our investigation of chemical communication in *Mastrus ridens* was motivated by the idea that the ecological function of sex pheromones depends on how males and females move through space. By mediating encounters between dispersing individuals, pheromonal communication can influence where mating occurs and therefore shape the mating system (Boulton et al. 2023, Hardy et al. 2023). This perspective provides the framework for the discussion below.

Males are markedly more mobile than females, taking off more readily and flying for longer. In response to female odor, they follow zig-zag or casting trajectories characteristic of pheromone-plume tracking and consistent with optomotor anemotaxis (Cardé 2021), allowing them to locate the odor source. They also hover around female-baited traps in the field, even under cloudy and windy conditions that appear unfavorable for flight. In rearing cages, they fly restlessly until encountering a female, which they then pursue on foot. Females, by contrast, are largely cryptic and pedestrian, remaining under substrates or along cage edges and usually escaping by running rather than flying.

This locomotor asymmetry reflects sex-specific roles: males engage in wide-ranging mate search guided by airborne cues, whereas females forage mainly on foot within confined microhabitats. Close-range host search relies on antennal contact and kairomonal cues from host cocoons (Jumean et al. 2005, Charles et al. 2019), yet females can also orient in flight toward host aggregations (Jumean et al. 2009). Because mating can occur while females are searching for hosts (Yáñez-Díaz et al. 2021), mate encounter in *M. ridens* is integrated into female foraging rather than occurring in a separate reproductive context (Fauvergue et al. 2008).

Dispersal combined with long-range sexual communication is expected to promote an outbreeding mating system. While dispersal is shaped by multiple selective pressures, it has long been regarded as a central mechanism of inbreeding avoidance (Bengtsson 1978, Pusey and Wolf 1996). More specifically, male-biased natal dispersal is predicted to evolve under the joint action of inbreeding depression and polygyny, in response to the costs of sibmating and male–male competition for access to females (Greenwood 1980, Pusey 1987, Li and Kokko 2019). Empirical studies in vertebrates are broadly consistent with this theoretical framework, and our results in *Mastrus ridens* point in the same direction. In this species, siblings emerge from the same host, yet sibmating is particularly costly because of single-locus complementary sex determination (sl-CSD; Retamal et al. 2016, Zaviezo et al. 2018). Males are polygynous, whereas females typically mate only once (Sandanayaka et al. 2011b), and despite repeated experiments, we found no behavioral evidence of sibmating avoidance. Thus, in *M. ridens*, outbreeding may rely primarily on dispersal and long-range mate location rather than on direct behavioral avoidance of relatives.

Comparable patterns occur in other hymenopterans with sl-CSD. In the solitary parasitoid *Venturia canescens*, emergence is spatially scattered, natal dispersal is weakly male-biased, and dispersing males orient toward both female and host odors, which act synergistically (Desouhant et al. 2005, Metzger et al. 2010b, Collet et al. 2020). Although females can discriminate sib from non-sib males, field populations show no genetic signature of sibmating avoidance (Metzger et al. 2010a, Collet et al. 2020), suggesting that dispersal and off-patch mate location is sufficient to limit inbreeding. Other species achieve outcrossing through different combinations of mechanisms. In *Bracon hebetor*, initial sexual unreceptivity, kin discrimination based on host-derived cues, and female dispersal to male rendezvous sites all reduce opportunities for sibmating (Antolin and Strand 1992, Ode et al. 1995, Guertin et al. 1996, Antolin et al. 2003). In the gregarious *Cotesia glomerata*, females show no direct sib avoidance but disperse from natal patches more often than males, apparently in response to interactions with related males (Gu and Dorn 2003, Ruf et al. 2010, Ruf et al. 2011).

Overall, the genetics and behavior of *M. ridens* support a male-biased dispersal strategy favored by strong inbreeding costs and polygyny, together with a mating system in which males actively locate females at a distance. This combination of traits contrasts with the classical LMC pattern of many parasitoid and fig wasps, characterized by local mating, frequent inbreeding, and female-biased dispersal. In such systems, haplodiploidy exposes recessive deleterious alleles to selection and thereby facilitates genetic purging (Henter 2003); males, sometimes apterous, remain on the natal patch (Hamilton 1979); and male–male competition is mitigated by strongly female-biased sex ratios (Hamilton 1967). In *Mastrus ridens*, the genetic burden imposed by sl-CSD may have turned a life history primed for local mating — siblings emerging together from the same host — into its evolutionary opposite: a system shaped by male dispersal, long-range mate finding via a volatile sex pheromone, and outbreeding.

## Conclusion

Our study reveals a sex-specific volatile communication system in *Mastrus ridens*, in which highly mobile males locate females at a distance and methyl 6-methylsalicylate emerges as a likely component of a multicomponent sex pheromone. By linking chemical communication to sex-specific movement and mate location, our results further suggest that pheromonal attraction contributes to an outbreeding mating system in a species strongly exposed to the genetic costs of inbreeding. This interpretation is supported by convergent evidence from laboratory behavior, field experiments, chemical analyses, and electrophysiology. Few studies of parasitoid sexual communication combine these levels of evidence within a single system. Earlier field and laboratory studies in *Lysiphlebus testaceipes* and *Leptopilina heterotoma* revealed how mate finding is embedded in parasitoid foraging and dispersal ecology (Fauvergue et al. 1999, Fauvergue et al. 2008) and in the latter, subsequent work resolved part of the chemical basis of this communication (Weiss et al. 2013). Comparable integrative efforts include *Campoletis chlorideae* (Guo et al. 2022) and *Venturia canescens* (Metzger et al. 2010b, Collet et al. 2016, Collet et al. 2020). In *M. ridens*, integrating these approaches reveals not only how males locate females, but how chemical communication may contribute to the evolution of the mating system itself.

## Acknowledgements

We thank the “Platform for Chemical Analyses in Ecology” (PACE), with the support of the LabEx CeMEB (Centre Méditerranéen pour l’Environnement et la Biodiversité, Montpellier, France) in particular Bruno Buatois for his invaluable advice. We also thank the INRAe unit “Plantes et Systèmes Horticoles” (PSH) for the insect supply and Marcel Thaon and Isabelle Le Goff for their great help with their husbandry.

